# CUT&Tag and DiBioCUT&Tag enable investigation of the AT-rich and dynamic epigenome of *Plasmodium falciparum* from low input samples

**DOI:** 10.1101/2024.06.24.600379

**Authors:** Jonas Gockel, Gala Ramón-Zamorano, Tobias Spielmann, Richárd Bártfai

## Abstract

Phenotypic variation between malaria parasites is one of the major contributors to the pathogens success and is regulated by differences in heterochromatin-mediated gene silencing. Currently, the heterochromatin landscape is mostly profiled utilising chromatin immunoprecipitation followed by next-generation sequencing (ChIP-seq). However this technique has drawbacks regarding AT-content-related artifacts and requires substantial material and time investment, severely limiting profiling of scarce sample types (e.g. field isolates). In order to facilitate assessments of epigenetic states in low-input samples, we adopted the epigenetic profiling technique Cleavage Under Targets and Tagmentation (CUT&Tag) to *Plasmodium falciparum* parasites. Performing the reaction with 100,000 or even only 10,000 nuclei yielded reproducible results coherent with bulk-ChIP-seq data while using significantly less material. We also optimised sample preparation, permitting the use of crude saponin lysates, which decreases sample loss due to inefficient nuclei isolation and increases versatility of the protocol. Finally, we developed DiBioCUT&Tag, a novel way of utilising dimerisation-induced recruitment of biotin ligases for signal amplification prior to anti-biotin CUT&Tag, which we successfully deployed to profile both heterochromatin occupancy and a dynamically chromatin-associated protein (BDP5). Methods described here hence provide substantially improved means for epigenetic profiling of (transiently) chromatin-associated proteins from low-input samples in the malaria parasite and beyond.

## INTRODUCTION

Epigenetic regulatory mechanisms influence cellular differentiation via activating or repressing expression of genes that impact cellular fate. Furthermore, they enable short-term adaptability of organisms next to longer-term adaptation due to DNA sequence alterations. Epigenetic regulation is mainly achieved by posttranscriptional modification of histones and consequent alteration on chromatin structure and accessibility via the effector proteins. A classical and conserved example is the formation of transcriptionally inactive heterochromatin via deposition of methyl marks on lysine 9 of histone H3 (H3K9me3) and consequent binding and oligomerisation of heterochromatin protein 1 (HP1)[1-5].

While epigenetic regulation is extensively studied in the context of vertebrate development, it is much less understood in eukaryotic pathogens, like *Plasmodium falciparum*, the causative agent of malaria. Nonetheless, accumulating evidence suggests that developmental transitions in the complex life cycle of these parasites, transmitting between human and arthropod hosts, are largely dependent on epigenetic regulation [6-8]. Furthermore, heterochromatin-mediated silencing, at the chromosome ends and in some chromosome internal islands, contributes to drug resistance, host-specific adaptation of invasion ligands and evasion of the immune system via antigenic variation [9-13]. Finally, given its essential function, epigenetic regulation is a potent target for drug development [14].

Previous studies investigating genome-wide distribution of epigenetic marks and chromatin-associated proteins in malaria parasites almost exclusively employed Chromatin Immunoprecipitation (ChIP) followed by next-generation sequencing [7, 15, 16]. ChIP relies on affinity-based purification of formaldehyde-crosslinked and sonicated chromatin fragments containing a transcription factor or epigenetic marks of interest targeted with a specific antibody. This technique combined with next-generation sequencing is generally regarded as the gold standard in the epigenomic field. While ChIP-seq can provide reliable results, it has some limitations [17]: i) given the inefficiency of immunoprecipation it requires millions of cells as input material; ii) chromatin fragmentation by sonication is batch sensitive; iii) formaldehyde fixation can influence antibody binding, cannot capture transient interactions and leads to biases for GC-rich sequences. These shortcomings are further exacerbated in *Plasmodium falciparum* owing to amplification and sequencing biases during analysis of its extremely AT-rich genome [18-20]. Collectively, while ChIP-seq has been instrumental in initial exploration of the *Plasmodium* epigenome, it does not allow for investigation of any samples with limited availability of material such as field isolates, mosquito and liver stage parasites, let alone individual parasites.

Novel epigenetic profiling techniques developed for model eukaryotes, such as CUT&Run [21] or CUT&Tag [22] are aiming to minimise these short comings of ChIP-seq. To perform a CUT&Tag experiment, nuclei are isolated and then bound to concanavalin A beads as a platform for subsequent incubation and wash steps. Epigenetic marks of interest are targeted with a specific antibody and incubated with the nuclei, followed by a secondary antibody to amplify the signal. These antibodies then direct a proteinA-Tn5 transposase fusion protein to the specified modifications, at which loci tagmentation of sequencing adapters is induced by addition of Mg2+ ions. All these processes occur within the nuclei, which are then lysed and their DNA extracted. Libraries can be prepared by PCR amplification of short fragments with two integrated adapter sequences, which following size selection are sequenced on a compatible NGS platform. Therefore, chromatin is not randomly fragmented by mechanical stress, but instead fragments are generated by integration of sequencing adapters, which are then specifically amplified. CUT&Tag is more efficient than ChIP-seq with significantly reduced background signal and can therefore be performed on limited input material and has even been used for single cell epigenetic profiling of histone marks [22-24].

In this work, we adapted and optimized the epigenetic profiling technique CUT&Tag to *Plasmodium falciparum* parasites. Our protocol is able to reliably and consistently profile heterochromatin landscape on both H3K9me3 and HP1 antibody targets. Furthermore, we show that CUT&Tag is both suitable for low-input down to 10.000 nuclei as well as crude whole parasite isolations. Lastly, we show the utility of a novel approach utilising dimerisation-induced recruitment of a biotin ligase (miniTurbo) via a chromatin associated protein (HP1 or BDP5) and CUT&Tag profiling of the corresponding chromatin regions via an α-Biotin antibody. This novel approach, we named DiBioCUT&Tag, enables profiling of transient chromatin binding events in the malaria parasite and beyond.

## MATERIAL AND METHODS

### *P. falciparum* cell culture

*P. falciparum* intraerythrocytic parasites were cultured at 37 °C under low oxygen conditions (3% O2, 4% CO2 and 93% N2) in human red blood cells at 5% hematocrit in RPMI 1640 medium supplemented with 0.2% NaHCO3 and 10% human serum. Wild type parasites were grown in the absence of antibiotics. For DiBioCUT&Tag, the HP1-2xFKBP-GFP-2A-NeoR / mCherry-FRB-miniTurbo and BDP5-2xFKBP-GFP-2A-NeoR / mCherry-FRB-miniTurbo strains were cultured in both the presence of Gentamicin G-418 sulfate (400 ug/mL, Invitrogen, ant-gn-5) and Blasticidin (0.4 ug/mL, Invitrogen, ant-bl) in modified RPMI with L-glutamine and without biotin and phenol red (US biological life sciences, R9002-01) supplemented with 200 µM Hypoxyxanthine (Merck, H9377) and 0.5% AlbuMAX ^TM^ II (Gibco, 11021037). Medium was furthermore supplemented with 250 nM Rapalog 1h before harvest and 50 µm Biotin (Invitrogen, B20656) 30 min before harvest. In order to achieve synchronicity, cultures were subjected to sorbitol-based lysis of remodeled RBCs [25].

### CUT&Tag and DiBioCUT&Tag: nuclei preparation

Parasite cultures were lightly crosslinked with 0.1% formaldehyde (Sigma, F8775), incubating for 2 min at 37 °C while shaking. Crosslinking was stopped by addition of glycine to 0.125 M final concentration. Samples were handled on ice from here onwards.

Cells were harvested by 440 g centrifugation for 8 min at 4 °C, and washed with ice-cold PBS. Centrifugation was repeated and the pellet was washed with PBS supplemented with 1x EDTA-free Protease Inhibitor (Roche, 04693132001). Parasites were extracted by adding saponin (0.05% total concentration) and incubating at room temperature (RT) for 10 min. Nuclei were isolated by carefully transferring the extracted parasite mixture on top of a 0.25 M to 0.1 M sucrose gradient in cell lysis buffer (10 mM Tris pH 8, 3 mM MgCl2, 0.2% NP-40, 1x EDTA free Protease inhibitor (Roche, 04693132001); 15 mL 0.25 M Sucrose and 17.5 mL 0.1M Sucrose for 50 mL tubes, 4 mL 0.25 M Sucrose and 6 mL 0.1 M Sucrose for 15 mL tubes) and centrifuging for 12 min, 3100 g, 4 °C with acceleration and deceleration set to 1 (Eppendorf 5910 Ri, Rotor S-4×400).

Supernatant was removed and nuclei washed in cell lysis buffer (10 min, 3500 g, 4 °C; Heraeus Fresco 21). Nuclei were counted in an automatic hemocytometer (BioRad, TC10 Automated Cell Counter) and were directly used for CUT&Tag. Optionally, nuclei were washed with cell lysis buffer containing 20% Glycerol and nuclei pellet was snap frozen in liquid nitrogen prior to storage at -80°C.

### CUT&Tag and DiBioCUT&Tag: parasite isolation

Parasite cultures were harvested by 440 g centrifugation for 8 min at RT and pellet was resuspended in 1 mL of 0.15% Saponin/PBS per 10 mL culture and incubated for 5 min on ice.

Samples were vortexed shortly every minute. Samples were washed three times with ice-cold PBS (3500 g centrifugation for 3 min at 4 °C; Heraeus Fresco 21). Intact parasites were counted in an automatic hemocytometer and were directly used for CUT&Tag. Optionally, parasites were washed with PBS containing 20% Glycerol and parasite pellets snap frozen in liquid nitrogen prior to storage at -80 °C.

### CUT&Tag and DiBioCUT&Tag: antibody incubation and tagmentation

Nuclei or whole parasite were resuspended in CUT&Tag wash buffer (20 mM HEPES pH 7.5, 150 mM NaCl, 0,5 mM Spermidine, 1x EDTA-free protease inhibitor) containing 0.1% Triton X-100 and permeabilized for 10 min on ice. Nuclei / parasites were pelleted by centrifugation (3500 g, 10 min, 4 °C) and resuspended in CUT&Tag wash buffer. Concanavalin A beads (Bangs Laboratories, BP531) were activated by resuspending into 10 volumes of Bead Binding Buffer (20 mM HEPES pH 7.5, 10 mM KCl, 1 mM CaCl2, 1 mM MnCl2), washed once on a magnetic rack and resuspended in the starting volume of bead slurry. Purified nuclei or parasites were bound to 10 µL beads per reaction (or 3 µL beads for low-input samples) by incubating for 10 min rotating at RT. The supernatant was removed and 50 µL antibody buffer (CUT&Tag wash buffer; 2 mM EDTA, 0.1% BSA) with primary antibody (0.25 µL polyclonal rabbit αHP1 [11]; 0.5 µg αH3K9me3, Abcam 8988; 0.5 µL αBiotin (Cell Signaling Technology # D5A7; for DiBioCUT&Tag) or 0.5 µg normal rabbit IgG (MERCK, # 12-370)) was added and incubated nutating over night at 4 °C. Unbound primary antibody was removed by washing once with 100 µL CUT&Tag wash buffer and samples were then incubated with secondary antibody (1.2 µg guinea pig anti-rabbit antibody, Antibodies-Online ABIN101961, in 100 µL CUT&Tag wash buffer, 1:100 dilution) for 1h on RT, nutating. The nuclei or parasites were washed on a magnetic stand twice with 100 µL CUT&Tag wash buffer and once with 100 µL CUT&Tag 300 Wash Buffer (20 mM HEPES pH 7.5, 300 mM NaCl, 0,5 mM Spermidine, 1x EDTA-free protease inhibitor). For all following wash steps CUT&Tag 300 wash buffer was used in order to quench potential affinity of protA-Tn5 to accessible chromatin regions. 2.5 µL commercial proteinA/G-Tn5 fusion protein (CUTANA™ pAG-Tn5 for CUT&Tag, Epicypher, 15-1017) was added in 50 µL CUT&Tag 300 wash buffer and incubated nutating for 1 h. Unbound proteinA/G-Tn5 fusion protein was removed by thrice washing with 100 µL CUT&Tag 300 wash buffer. The supernatant was removed and the nuclei or parasites were resuspended in 200 µL freshly prepared tagmentation buffer (CUT&Tag 300 wash buffer, 10 mM MgCl2). To perform tagmentation, samples were incubated in a PCR thermocycler (BioRad, T100) at 37 °C for 1h. Tagmentation was stopped and nuclei or parasite lysis was facilitated by addition of 10 µL of 0.5M EDTA pH8, 3 µL of 10% SDS and 1 µL of 50 mg/mL proteinase K. Samples were briefly vortexed and then incubated at 55 °C for 1 h. DNA fragments were extracted utilizing the DNA Clean & Concentrator -5 kit (Zymogen, D4014) following manufactures instructions. DNA was eluted from the column with 26 µL of prewarmed elution buffer and DNA concentrations were assessed with Qubit dsDNA High Sensitivity Assay kit (Invitrogen, Q33231).

### CUT&Tag and DiBioCUT&Tag: library preparation

Maximal 50 ng of extracted DNA from CUT&Tag experiments were amplified with unique combinations of i5 and i7 barcoded primers [26], enabling tracing the DNA fragments originating from separate experiments in the sequencing data. PCRs were performed in a total reaction volume of 50 µL using Kapa HiFi polymerase (use non-hotstart version for gap filling; Roche, KK2102) in a thermocycler with the following program: 58 °C for 5 min, 62 °C for 5 min (gap filling), 98 °C for 2 min, 12-16 cycles of 98 °C for 20 second and 62 °C for 10 seconds, 62 °C for 1 min and hold at 4 °C. Post-PCR DNA cleanup was performed by adding 50 µL (1x volume) of AMPure XP bead slurry (Beckman Coulter, A63882) and incubating for 10 min at RT, washing twice with 80% EtOH on a magnetic rack, and eluting in 16.5 µL of 10 mM Tris-HCl pH 8 for 5 min at RT. DNA concentrations of libraries were assessed with Qubit dsDNA High Sensitivity Assay kit (Invitrogen, Q33231) and library fragment size distribution was accessed by microfluidic gel electrophoresis (Agilent 2100 Bioanalyser) with the corresponding High Sensitivity DNA Kit (Agilent, 5067-4626). Concentrations from successful libraries were between 0.2 µg/mL to 10 µg/mL, showing hints of a nucleosomal profile with average fragment length from 500-700 bp indicative of effective tagmentase activity. Sequencing was performed using an Illumina NextSeq 2000 instrument for 3-10 M reads per CUT&Tag sample; 59bp paired-end reads were generated.

### Sequencing Data analysis

Sequencing reads were mapped against the reference genome PlasmoDB v26 3D7 using bowtie2 (v2.5.2) with paired-end mapping for CUT&Tag data and single-end mapping in case of the NF54 HP1 ChIP-seq dataset [7]. Duplicates in the ChIP-seq datasets were removed with picard (v3.1.0). Duplicate removal was skipped for CUT&Tag datasets as duplicates may result due to the affinity of Tn5 to certain sequences as well as accessibility of certain regions leading to fragments with the same start and end locations [27]. Downstream analysis was furthermore not influenced by duplicate removal in pilot analysis of the CUT&Tag data sets.

Reads were filtered by mapping quality >= 30 and mitrochondrial as well as apicoplast reads were removed with samtools (v1.18). BigWig files normalized to read per million per kilobase (RPMK) were created using deeptools (v3.5.4) for binning of sequencing data into either 500 bp windows for log2 ratio track calculations or 2000 bp windows for correlation analysis. BedGraph files were generated for visualisation purposes on the UCSC genome browser with bedtools (v2.31.0) normalized to library size. A detailed & customisable script can be found on our github page https://github.com/bartfai-lab/DiBio-CUTnTag-Analysis.

Log2 ratio tracks were generated by running the multiBigwigSummary command from deeptools (v3.5.4) using 500 bp bins on both sample and control bigwig. A pseudocount of 1 was added to all values to prevent divisions by 0, and log2 values were calculated in bash and appended to a new bedGraph file for visualisation on UCSC genome browser.

### Correlation Analysis

Average enrichment scores of previously generated bigwig files for read count or log2 ratio tracks were calculated with multiBigwigSummary from the deeptools package (v3.5.4) on 2000 bp bins. Datasets were imported into R and zero values in the log2 ratio datasets were removed as these are most likely artefacts from adding pseudocounts to non-mappable or not-sequenced genomic regions. If different antibodies were compared, average enrichment scores were filtered for a minimum enrichment score prior to quantitative correlation analysis to remove variations from the background signal in non-background corrected tracks. Scatterplots were generated utilizing the ggplot2 package (v3.4.4), trendlines and R^2^ values were calculated using the ggpmisc (v0.5.4-1) package in R (v4.3.1).

### Modeling of background distribution and identification of regions with significant enrichment

Normal distribution was modeled on 2000 bp binned average enrichment score files previously obtained with the multiBigwigSummary command from the deeptools package (v3.5.4). Log10 values for the enrichment scores were calculated in order to enable proper visualisation and the normalmixEM function from the mixtools package (v4.3.1) was used to model the normal distribution of the background signal in R. Regions were considered significantly enriched if they showed higher enrichment than the top 0.005% of the background distribution (i.e. 0.5% false discovery rate).

### Peak identification of BDP5 DiBioCUT&Tag and expression analysis

Peaks were identified using macs2 (v2.2.9.1) using the IgG CUT&Tag background track as a control and a q-value of 0.005. Any peaks falling in telomeric regions before the first annotated var gene were removed with bedtools intersect (v2.31.0). For each peak, the closest ATG was identified using bedtools closest (v2.31.0) and peak profiles were generated using the deeptools package (v3.5.4). For expression analysis, peaks were intersected with the 5’ end of genes (-1000bp from ATG and 500bp into the coding body). RNA expression values of affected genes at different timepoints were obtained from Toenhake et al, 2018 RNAseq dataset [28]. Mean expression values of all affected genes were calculated and plotted using the ggplot2 package (v3.4.4). To generate heatmaps, expression values per gene were scaled to 1 and heatmaps were generated in R using the heatmap package (v1.0.12).

## RESULTS & DISCUSSION

### CUT&Tag enables efficient heterochromatin profiling in even the extremely AT-rich genome of *P. falciparum*

To implement CUT&Tag for *P. falciparum* parasites, we isolated and permeabilized nuclei from trophozoite and schizont stages. We followed the basic principle of the technique described by Kaya-Okur [22, 29] (Figure 1A), optimizing both nuclei isolation, permeabilisation and PCR protocols to *P. falciparum* cells (see Material and Methods for details). Genome wide occupancy profiles of both HP1 and H3K9me3 CUT&Tag showed a heterochromatin landscape very similar to ChIP-seq profiles (Figure 1B). To test for any technical biases towards GC-rich heterochromatic or other genomic regions influencing CUT&Tag results, we also performed CUT&Tag with a non-specific IgG antibody. The resulting genome occupancy tracks showed an almost flat background profile (Figure 1B, light blue), indicating the lack of substantial biases. Utilising the IgG read count as background, we also corrected the HP1 CUT&Tag read counts into a log2 ratio track providing an even more accurate measure of heterochromatin occupancy (Figure 1B/C). We next performed quantitative analysis in 2000 bp windows throughout the genome to compare HP1 CUT&Tag and ChIP-seq (Figure 1C) as well as HP1 and H3K9me3 CUT&Tag datasets (Figure 1D), both of which showed strong positive correlation demonstrating accuracy and reliability of the obtained heterochromatin profiles. To verify the sensitivity of our protocol, we examined heterochromatin differences between two different *P. falciparum* strains, previously reported by Fraschka et al. [7]. We observed the expected strain specific differences between NF54 and 3D7 (F12), including an extension of the heterochromatic domain on the proximal end of chromosome 12 (Figure 1E). These observations demonstrate the utility and accuracy of CUT&Tag for heterochromatin profiling even on an extremely AT-rich genome as that of the malaria parasite, *P. falciparum*.

**Figure 1:**
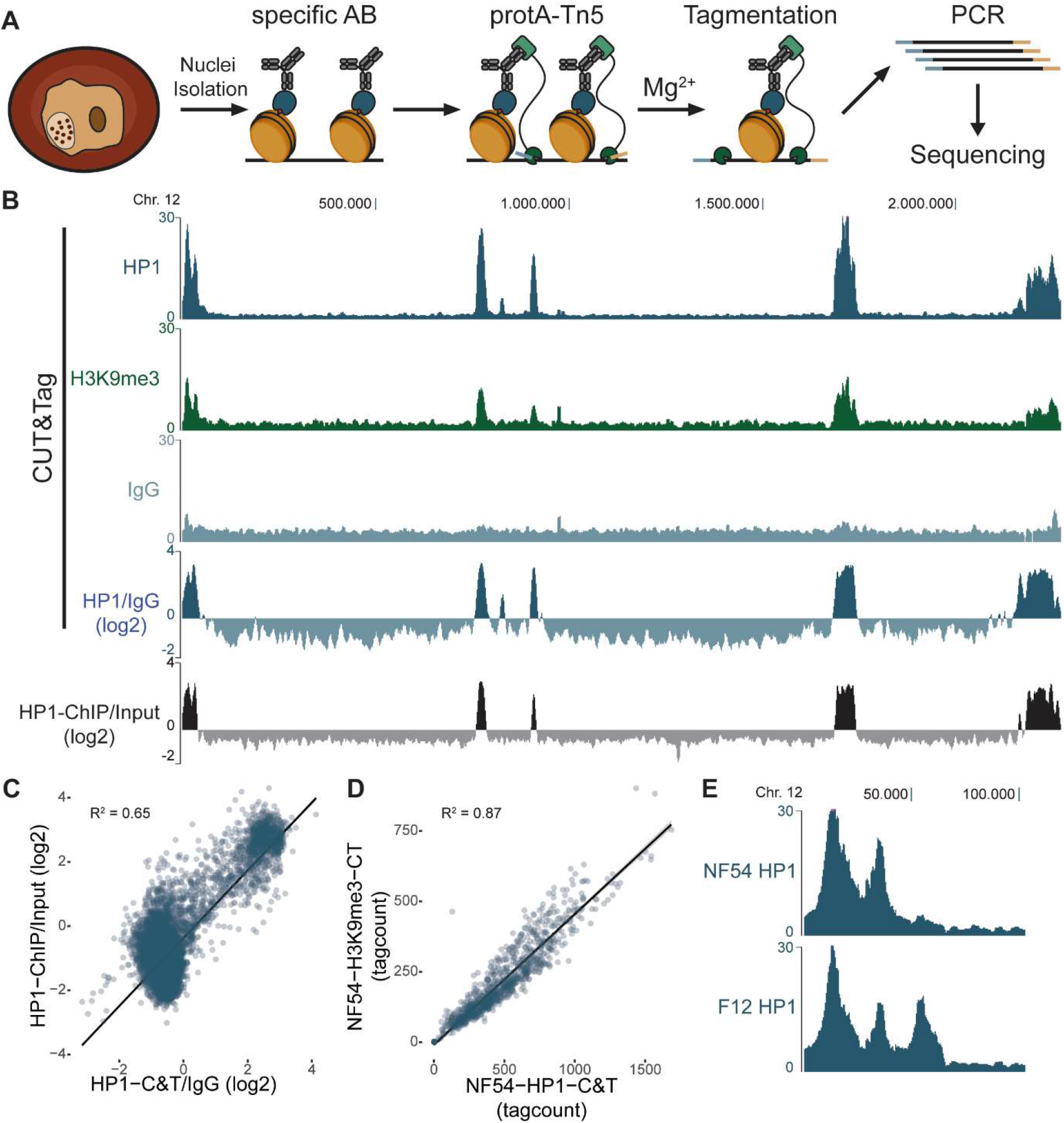
CUT&Tag of HP1 and H3K9me3 in Plasmodium falciparum provides accurate means to genome-wide heterochromatin profiling. **A)** The main steps of CUT&Tag. Nuclei are isolated from parasites and incubated with a specific antibody (AB) against the target of interest. This AB directs a proteinA-Tn5 transposase fusion protein to specific chromatin regions. Transposition of sequencing adaptors (tagmentation) is induced by addition of Mg2+ ions, and the resulting DNA fragments are amplified by PCR and then subjected to massive parallel sequencing. **B)** Chromosome-wide CUT&Tag profiles for HP1 (blue), H3K9me3 (green) and control IgG (normalized read count or log2 ratio tracks, light blue) as well as a HP1 ChIP-seq (ChIP/input, black). Log2 ratio tracks have been calculated in 500bp windows. **C)** Scatter plot displaying correlation between HP1 ChIP-seq and CUT&Tag log2 ratio tracks in 2000bp windows. **D)** Scatter plot displaying correlation between HP1 and H3K9me3 CUT&Tag read counts in 2000bp windows. Euchromatic regions (tag counts < 150) are not shown. **E)** HP1 CUT&Tag normalized tag count tracks at the distal end of chromosome 12 in two different *P. falciparum* strains (NF54 and F12) highlighting strain-specific differences in heterochromatin occupancy as earlier described by Fraschka et al [7].

### CUT&Tag can be scaled down to 10,000 nuclei without losing heterochromatin calling efficiency

One of the major advantages of CUT&Tag over ChIP-seq is its potential to scale down input material and profile heterochromatin in sparce sample types. To test whether CUT&Tag is applicable for these low-input conditions, we performed HP1 CUT&Tag on as little as 10.000 nuclei. Genome wide occupancy profiles between 100.000 and 10.000 nuclei showed very similar heterochromatin landscapes (Figure 2A), with a slight decrease in signal to noise ratio in the 10.000 nuclei sample. Background and signal however remained still clearly distinguishable (Figure 2B,C) and enabled efficient heterochromatin calling (Figure 2A, black boxes) with 0.5% false discovery rate (FDR) based on a modeled normal distribution of background signal (Figure 2B,C orange area). Furthermore, quantitative analysis showed high positive correlation between low and regular input reactions (Figure 2D). Therefore, CUT&Tag enables heterochromatin profiling from as little as 10.000 and possibly even less nuclei.

**Figure 2:**
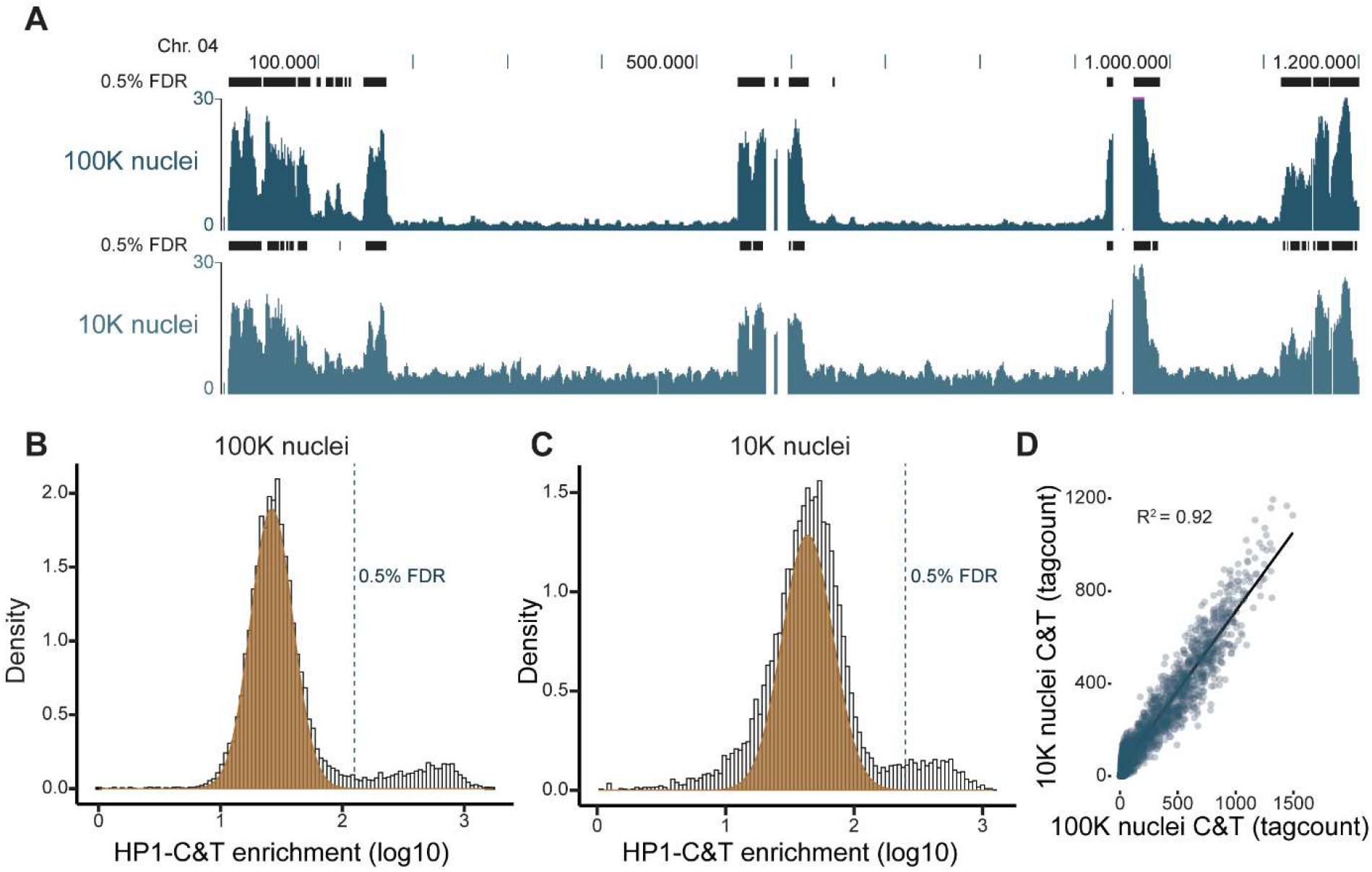
CUT&Tag enables identification of heterochromatic regions also from low input samples. **A)** Chromosome wide CUT&Tag profiles for HP1 using standard input (100,000 nuclei, dark blue) as well as low input (10,000 nuclei, light blue). Black bars represent 2000bp windows with signal significantly higher than background (as determined in Figure 2 B/C with a false discovery rate of 0.5%). **B)** Histogram depicting read count distribution in 2000bp windows for HP1 CUT&Tag using 100,000 nuclei. Normal distribution model (orange) for background signal and 0.5% FDR cutoff for heterochromatin calling (dotted line) are indicated. **C)** Histogram depicting read count distribution in 2000bp windows for HP1 CUT&Tag using 10,000 nuclei. Normal distribution model (orange) for background signal and 0.5% FDR cutoff for heterochromatin calling (dotted line) are indicated. **D)** Scatter plot displaying correlation between 100,000 and 10,000 nuclei samples in 2000bp windows.

### Nuclei isolation is not essential for CUT&Tag, enabling more efficient sample processing from intact parasites

The process of isolating nuclei is both laborious and leads to substantial sample loss. Furthermore, direct processing of the samples for CUT&Tag (e.g. endemic settings) is not always possible and hence storage is desirable. To optimize our CUT&Tag protocol, we pursued the use of intact isolated parasites (either directly or after frozen storage) as an input material instead of nuclei (Figure 3A). Parasites were released from infected cells by saponin mediated lysis of the red blood cells and were then either used directly as CUT&Tag input or snap frozen for storage (Figure 3A). Importantly, neither skipping the nuclei isolation step (Figure 3B, Sup. Figure 1A) nor freezing (Figure 3C, Sup.Figure 1B) impaired heterochromatin landscape profiling. These results show that we can generate reliable results when using frozen parasite isolates for CUT&Tag experiments and in turn minimize sample preparation time and improve the efficiency and flexibility of these experiments.

**Figure 3:**
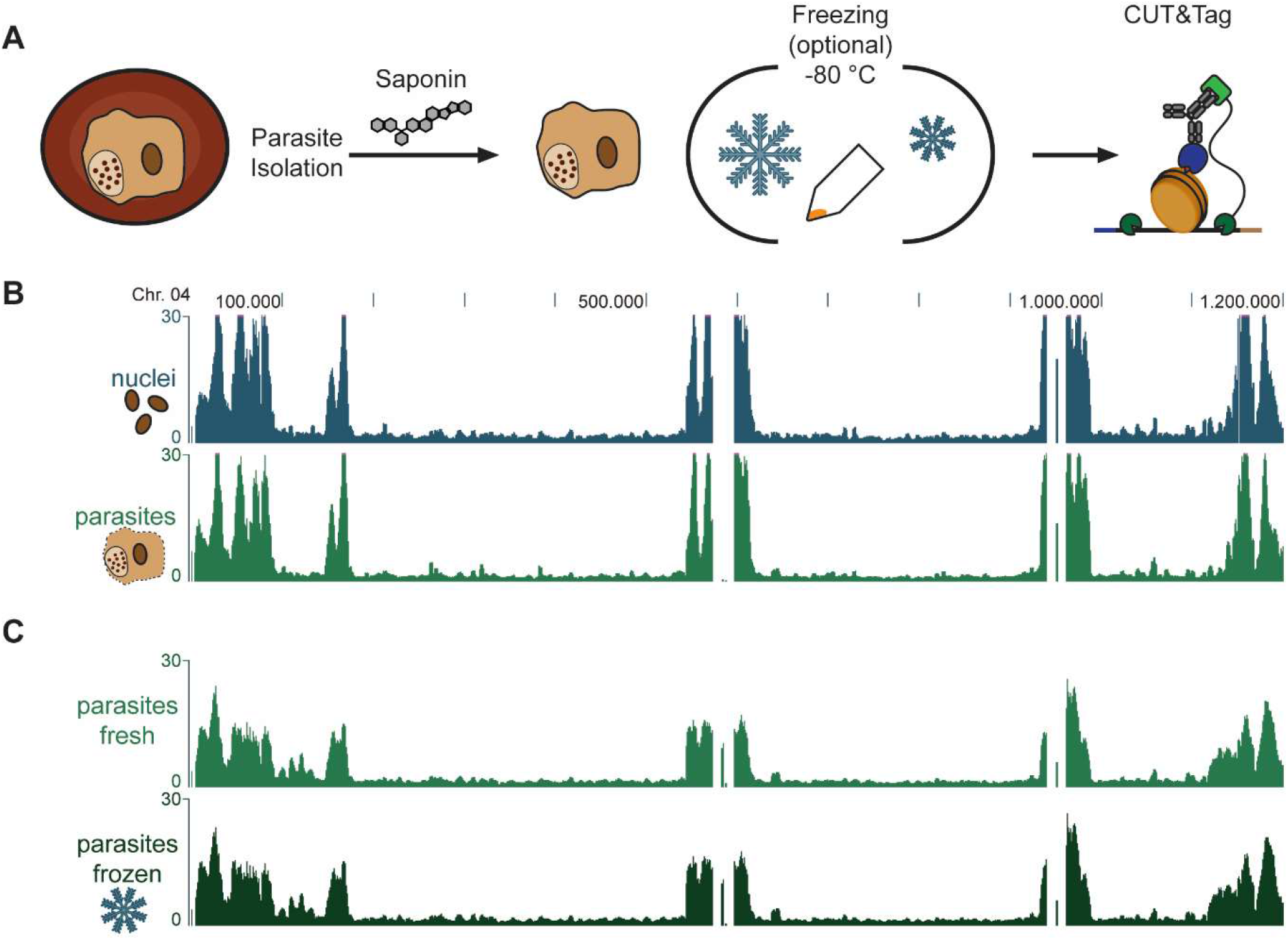
CUT&Tag can also be performed on frozen and/or intact parasites without isolation of nuclei. **A)** Schematics of parasite isolation for CUT&Tag. Parasites are isolated by saponin-mediated lysis of the red blood cell and either used directly for CUT&Tag or can be snap frozen in liquid nitrogen and stored at -80°C. **B)** Chromosome-wide HP1 CUT&Tag profiles using 1 million isolated nuclei (blue) and 1 million parasites (green). **C)** Chromosome-wide HP1 CUT&Tag profiles using 100K isolated parasites directly (fresh, light green) or after freezing (frozen, dark green).

The process of isolating nuclei is both laborious and leads to substantial sample loss. Furthermore, direct processing of the samples for CUT&Tag (e.g. endemic settings) is not always possible and hence storage is desirable. To optimize our CUT&Tag protocol, we pursued the use of intact isolated parasites (either directly or after frozen storage) as an input material instead of nuclei (Figure 3A). Parasites were released from infected cells by saponin mediated lysis of the red blood cells and were then either used directly as CUT&Tag input or snap frozen for storage (Figure 3A). Importantly, neither skipping the nuclei isolation step (Figure 3B, Sup. Figure 1A) nor freezing (Figure 3C, Sup. Figure 1B) impaired heterochromatin landscape profiling. These results show that we can generate reliable results when using frozen parasite isolates for CUT&Tag experiments and in turn minimize sample preparation time and improve the efficiency and flexibility of these experiments.

### DiBioCUT&Tag is a novel dimerisation-induced, proximity-labelling-based approach for epigenetic profiling

One of the main limitations of CUT&Tag is the difficulty of applying it to profiling of temporarily chromatin-associated factors (e.g. transcription factors, chromatin modifying enzymes or reader proteins). In order to overcome this limitation, we included an additional step of conditionally biotinylating strongly chromatin-associated proteins (e.g. histones) in the vicinity of target protein and performing CUT&Tag with an α-biotin antibody (Figure 4B). This, in principle, should lead to (stage-specific) accumulation of the signal over time and amplification of weak signals. As a proof of principle, we use a parasite line in which HP1 is FKBP tagged [30] and transfected it with a plasmid carrying a FRB - miniTurbo biotin ligase fusion protein (Figure 4A). Upon the addition of rapalog, FKBP and FRB dimerise and miniTurbo is recruited to HP1 occupied regions, where resident histones are then biotinylated (Figure 4B). Targeting these biotinylation events with an α-biotin antibody in a CUT&Tag reaction led to reliable heterochromatin profiling (Figure 4C). This demonstrates that DiBioCUT&Tag can be used for profiling of heterochromatin and most likely can be expanded to other chromatin associated proteins.

**Figure 4:**
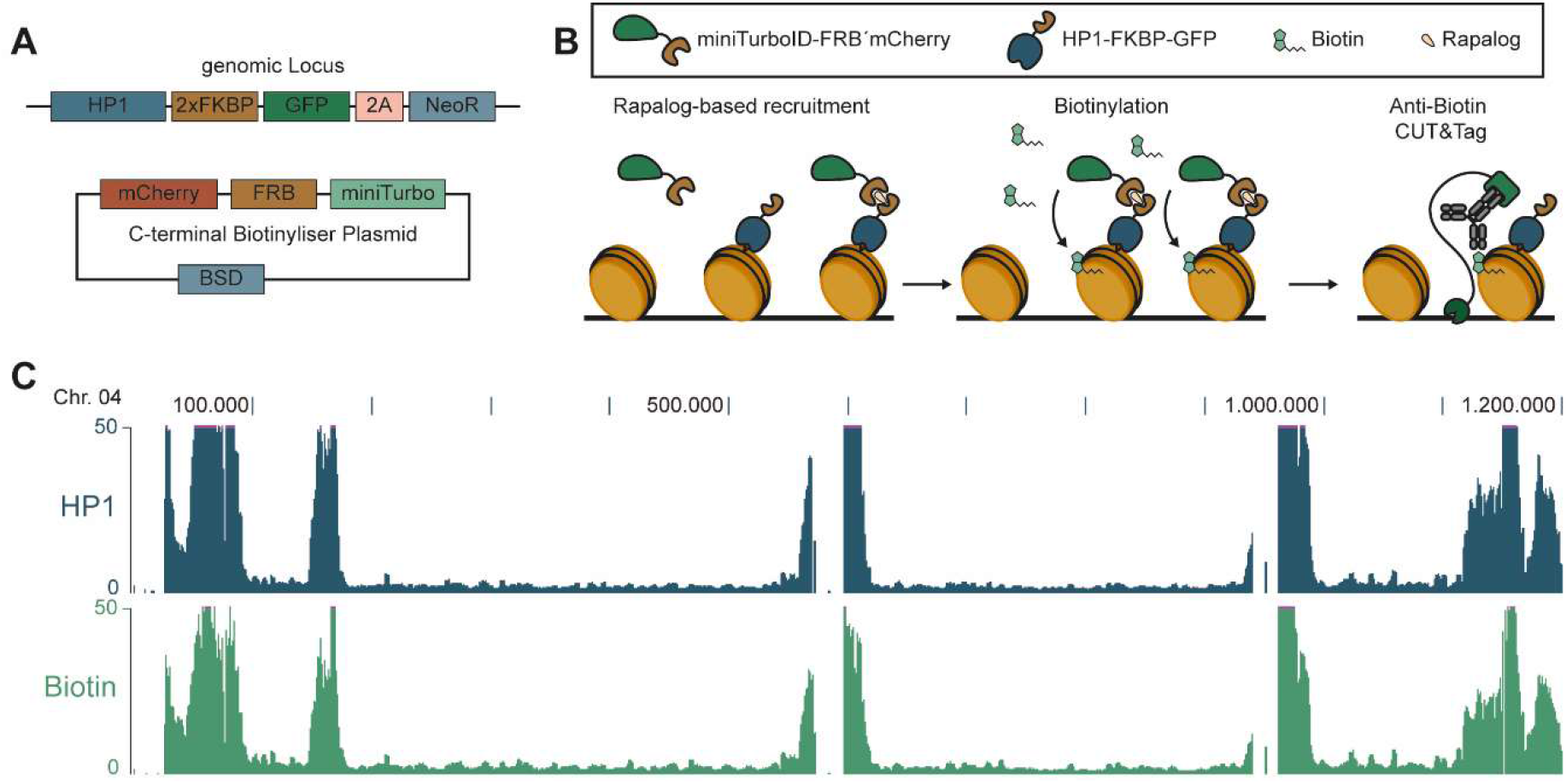
BioCUT&Tag is a proximity-labeling-based amplification for epigenetic profiling. **A)** Drawing depicting endogenous tagging of the HP1 locus with 2xFKBP-GFP [30] and an episomal plasmid carrying mCherry-FRB-miniTURBO construct. **B)** Schematic of BioCUT&Tag. The biotynyliser, miniTurbo-FRB-mCherry is recruited to the target protein, HP1-FKBP-GFP by addition of rapalog. In presence of biotin, chromatin-associated proteins in the vicinity of HP1 are biotinylated. These biotin molecules are then targeted with a specific anti-biotin antibody in the subsequent CUT&Tag reaction. **C)** Chromosome-wide read occupancy tracks from HP1 (blue) and biotin CUT&Tag (green).

### BDP5 associates with promotor regions of stage-specifically expressed genes

To prove the utility of the DiBioCUT&Tag approach for profiling of temporarily chromatin associated proteins, we applied it to characterise genome-wide binding of BDP5. Previous attempts to identify binding sites of this factor with ChIP-seq in our group have been unsuccessful, likely due to low residence time of this factor on its targets. To capture this dynamic occupancy we profiled “biotin footprints” left behind by BDP5-FKBP + FRB-TurboID dimers at two different parasite stages (trophozoites, 24-32 hpi and schizonts, 32-40 hpi) following short rapalog/biotin treatment. DiBioCUT&Tag gives rise to clear local enrichments and distinct peak profiles in both samples (Figure 5A), which could not be observed in generally flat and noisy “profile” obtained in absence of rapalog/biotin (Sup. Figure 2). Using a peak calling algorithm we identified 115 and 143 high-confidence peaks in the 24-32 hpi and 32-40 hpi sample, respectively. Most of these peaks are located in the 5’ regulatory regions of genes as demonstrated by clear accumulation of the average DiBioCUT&Tag signal upstream of the ATG (Figure 5B,C, brown lines) in comparison to the control (no Rapalog, no biotin; Figure 5B,C, blue lines). Genes with a BDP5 peak in their promotor region are mainly expressed in the corresponding parasite stage (Figure 5D, Sup. Figure 3A,B), indicating a correlation between BDP5-binding and gene expression. Therefore, DiBioCUT&Tag enables genome-wide profiling of transiently chromatin-associated proteins and further corroborates the previously predicted function of the BDP5-containing TAF1 complex in transcription initiation [31].

**Figure 5:**
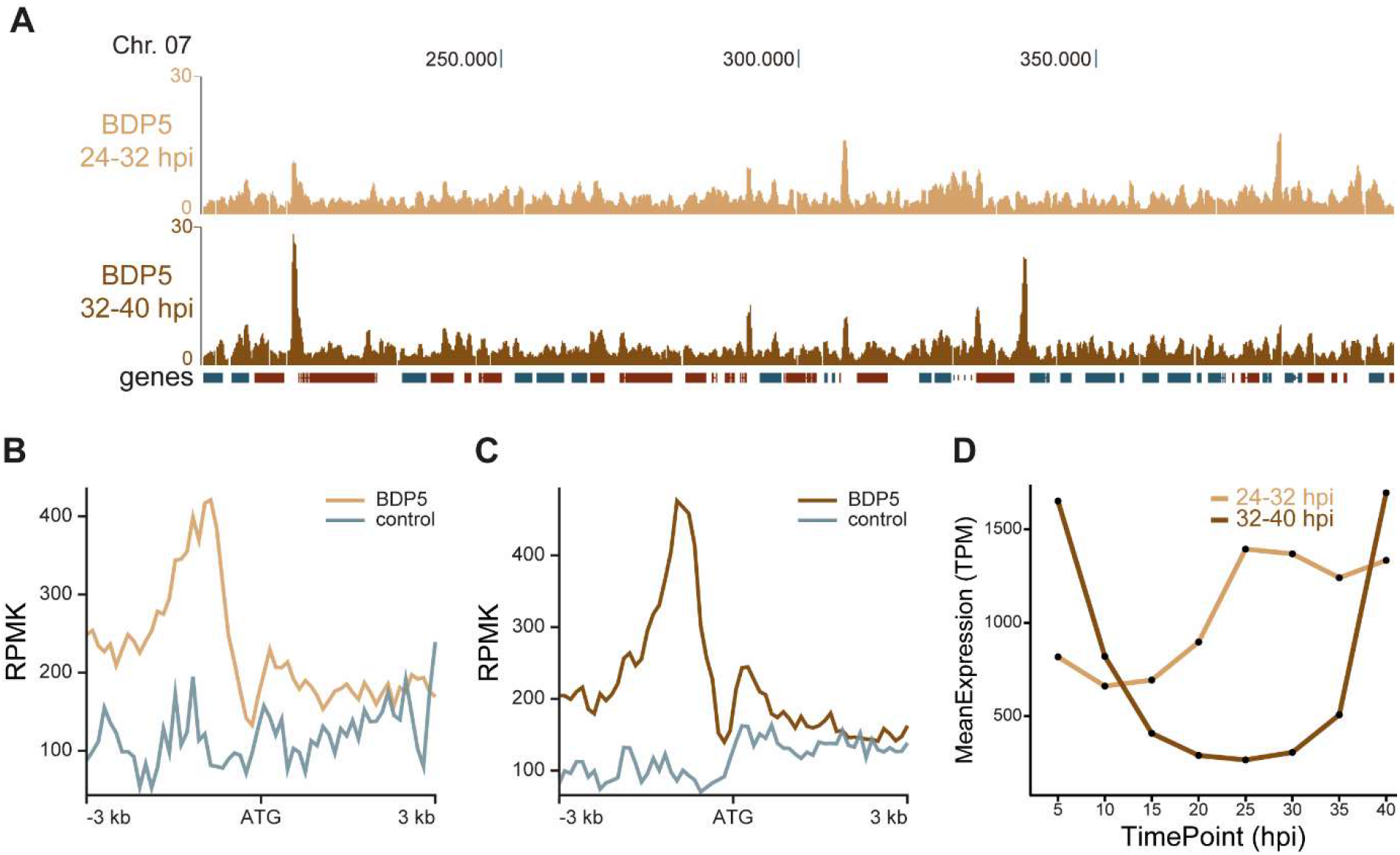
BDP5 dynamically associates with the promotor region of stage-specifically expressed genes. **A)** Read occupancy tracks of BDP5 DiBioCUT&Tag for trophozoite (24-32 hpi, orange) and schizont (32-40 hpi, brown) stage parasites. **B)** Peak profiles of 24-32 hpi BDP5 DiBioCUT&Tag (orange) and control (blue; no biotin and no rapalog) in relation to the ATG. **C)** Peak profiles of 32-40 hpi BDP5 DiBioCUT&Tag (brown) and control (blue; no biotin and no rapalog) in relation to the ATG. **D)**Mean expression of genes (transcripts per kilobase million) bound by BDP5 up to 1000 bp before and 500 bp after ATG at different times of the life cycle (24-32 and 32-40 hpi) as defined by Toenhake et al 2018 [28]. Detailed heatmaps can be found in Sup. Figure 3A,B.

## Conclusions

In this study, we show that CUT&Tag is a robust technique to reliably profile heterochromatic regions even in the extreme AT-rich genome of *Plasmodium falciparum*. Limiting input material to 10.000 nuclei still led to reproducible results with clear separation of euchromatic and heterochromatic regions. Furthermore, fresh and frozen intact parasite isolations provided very similar and reliable heterochromatin readouts and hence can be used to minimize sample loss during processing. Avoiding purification of nuclei and the capacity to store frozen samples also increases the potential range of sample sources (clinical or endemic settings) and simplifies as well as reduced the time of the protocol. CUT&RUN [32, 33] and CUT&Tag [34] have recently been used for profiling of histone modifications in *P. falciparum* parasites. However, their utility to low input samples and profiling of transient chromatin-binders was not addressed. Accordingly, the developments described here open new avenues towards investigation of scarce sample types, such as field isolates or mosquito and liver stages of parasite development. Furthermore, CUT&Tag has been proven to be applicable to the single cell level [22-24], which in case of *Plasmodia* has the potential to revolutionize the exploration of epigenetic variation between individual parasites that may underly developmental decisions or virulence factor expression on the individual cell level. Based on our experiments (Figure 2), however, we predict a further decline in signal to noise ratio and very sparce data at the single parasite level. Therefore, single haploid parasite CUT&Tag will most likely only be applicable to stable and broadly distributed epigenetic features, such as heterochromatin. However, DiBioCUT&Tag might sufficiently amplify signals of other chromatin bound factors to enable their profiling on single cell level.

Despite multiple advantages, CUT&Tag, similar to CUT&RUN and ChIP-seq depends on the availability of specific and compatible antibodies. Furthermore, high salt concentrations necessary to quench unspecific tagmentation events lead to inefficient capturing of transient interactions, such as binding of transcription factors and effector proteins [29]. To overcome these limitations we developed a novel approach where strongly chromatin associated proteins (e.g. histones) in the vicinity of the binding site are biotinylated and later profiled by anti-biotin CUT&Tag (DiBioCUT&Tag). CUT&Tag of the biotinylated chromatin enables the use of a standard α-biotin antibody [35], but requires genetic modification of the target proteins. Furthermore, direct coupling of the target protein and the biotin ligase can lead to background biotinylation and could prevent temporal resolution of dynamic binding patterns. Hence here we made the recruitment of the biotin ligase conditional via the FKBP-FRB interaction [36] and demonstrate feasibility of this approach by successful profiling HP1 as well as BDP5.

In summary, we present CUT&Tag and DiBioCUT&Tag as a reliable, cost and time effective alternative to ChIP-seq epigenetic profiling in *Plasmodium falciparum*. These advances will be imperative to analyse sparce sample types during the parasite life cycle, deepening our understanding of this deadly parasite as well as potentially revealing new epigenetic drug targets.

## DATA AVAILABILITY

All raw and processed sequencing data has been submitted to Gene Expression Omnibus (GEO) under reference number GSE270104. Code used for CUT&Tag data processing and visualization is available at GitHub https://github.com/bartfai-lab/DiBio-CUTnTag-Analysis.

## AUTHOR CONTRIBUTIONS

Jonas Gockel: Conceptualisation, Investigation, Methodology, Formal analysis, Visualisation, Writing—original draft. Gala Ramón-Zamorano: Resources, Investigation, Writing—review & editing. Tobias Spielmann: Resources, Funding Acquisition, Writing—review & editing. Richárd Bártfai: Conceptualisation, Funding Acquisition, Supervision, Writing—original draft.

## ACKNOWLEDGEMENTS

The authors thank Till Voss (Swiss TPH, Basel) for sharing the α-PfHP1 antibody with us and his advice throughout this project.

## Supplementary Figures

**Supplementary Figure 1:**
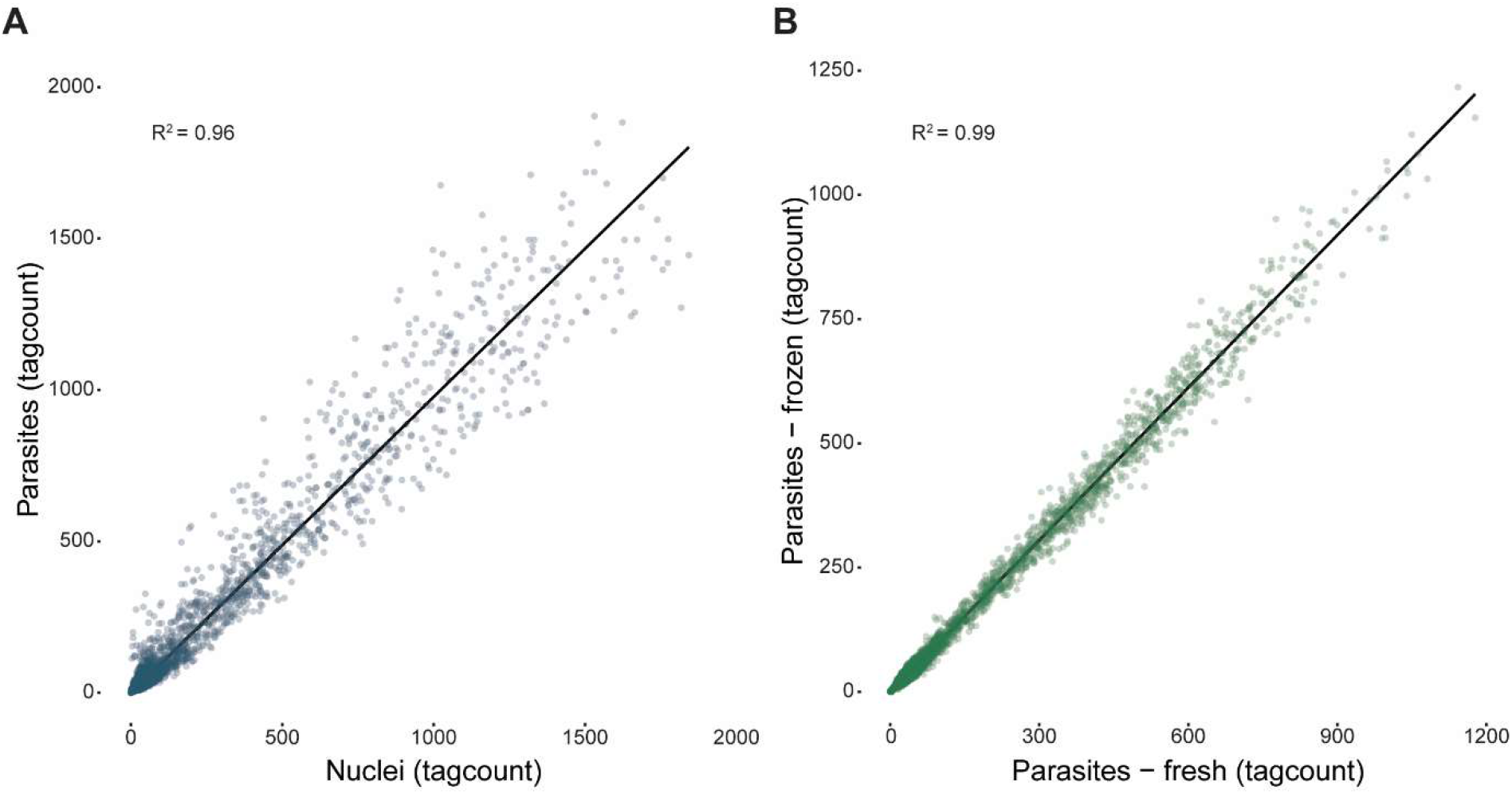
Scatter plots showing correlation of different parasite isolates. **A)** Scatter plot displaying correlation between isolated parasites and isolated nuclei as input material for CUT&Tag **B)** Scatter plot displaying correlation between frozen isolated and freshly isolated parasites as input material for CUT&Tag

**Supplementary Figure 2:**
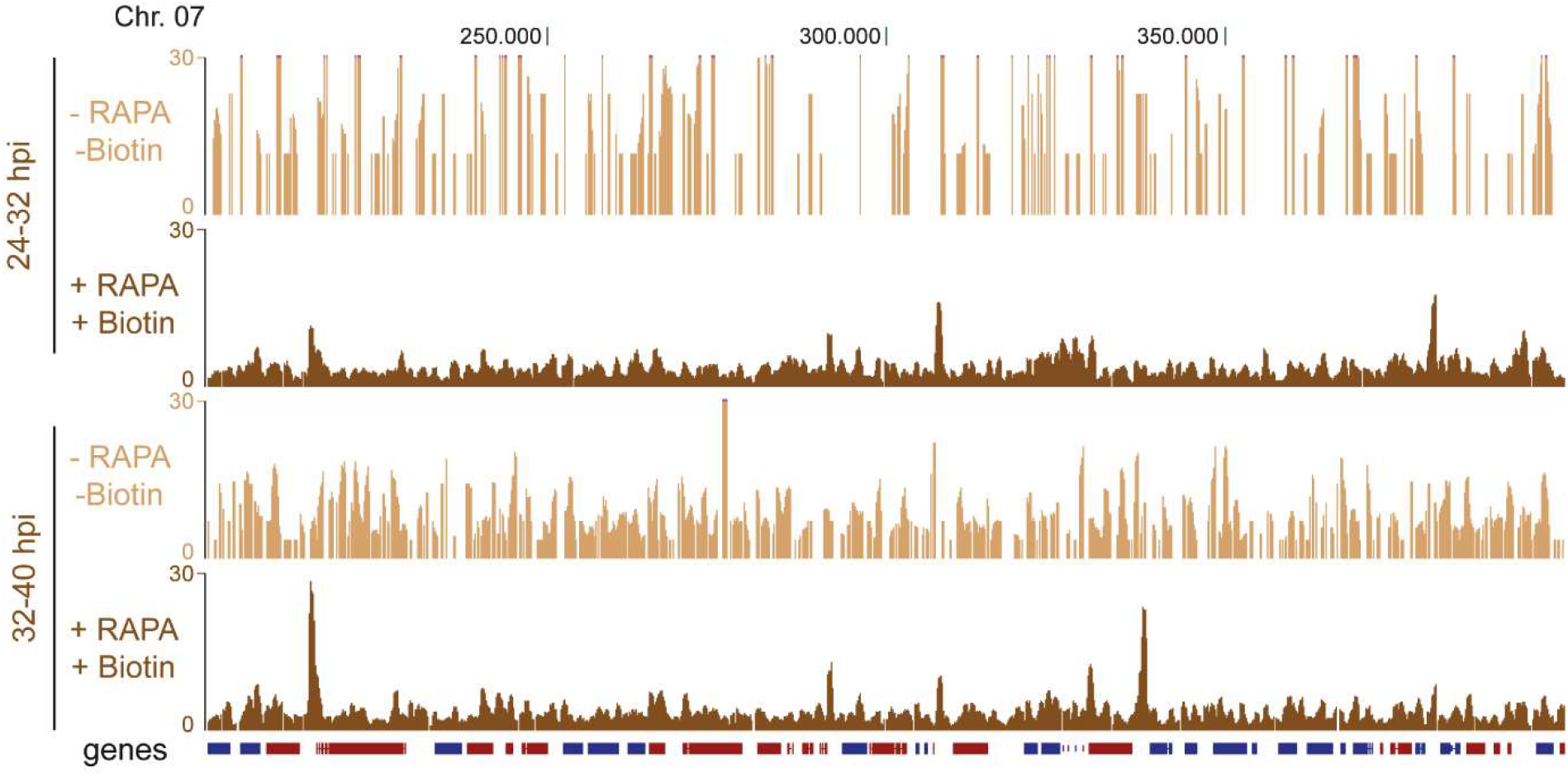
BDP5 read occupancy tracks with controls. Read occupancy tracks of BDP5 DiBio-CUT&Tag for trophozoite (24-32 hpi) and schizong (32-40 hpi) stages including respective controls without rapalog and biotin (light brown).

**Supplementary Figure 3:**
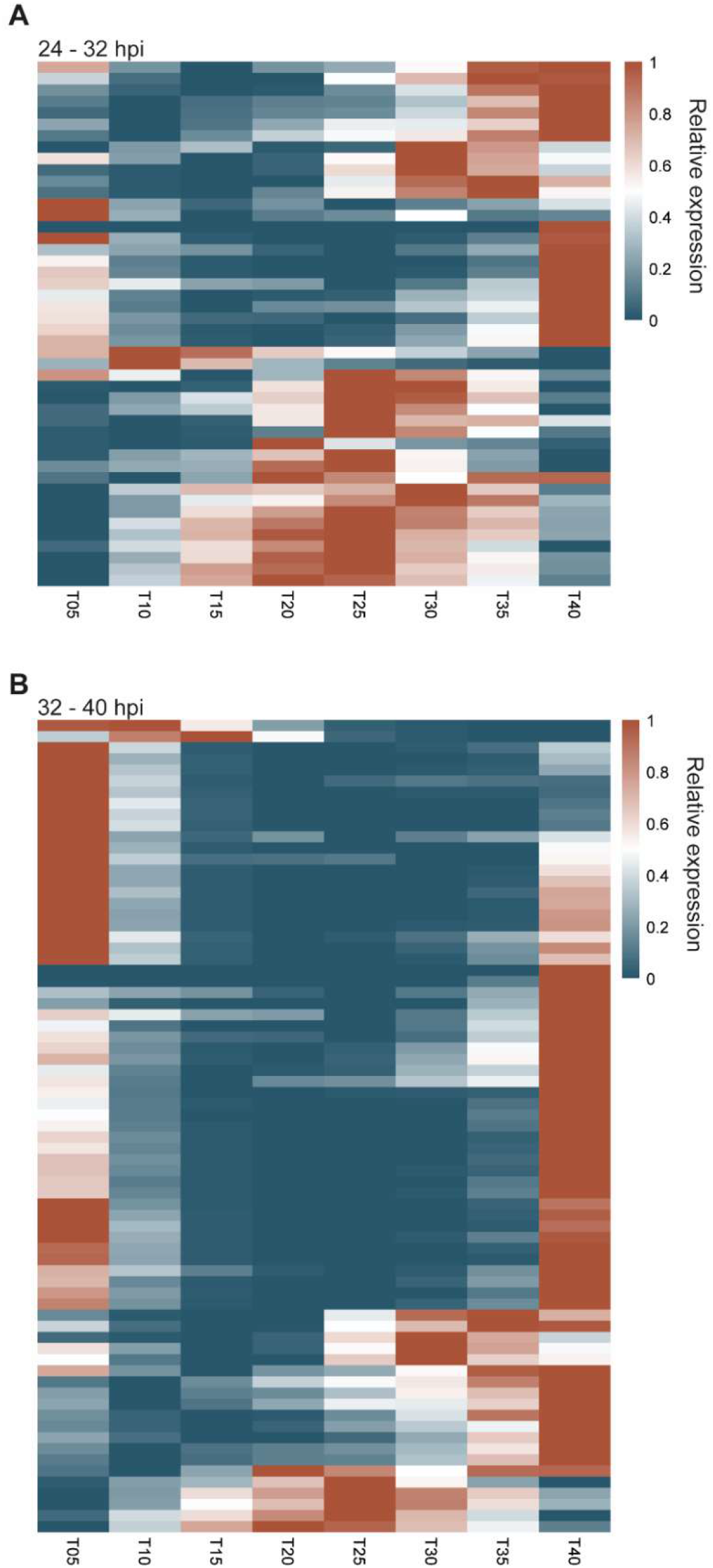
Heatmaps BDP5 affected genes during lifecycle. Heatmaps showing relative RNA expression during the asexual life cycle for genes which have BDP5 binding 1000 bp before and 500 bp after their ATG for trophozoites (**A**, 24-32 hpi) and schizonts (**B**, 32-40 hpi). RNAseq data was taken from Toenhake et. al, 2018.

